# A robust protocol for the systematic collection and expansion of cells from ER^+^ breast cancer tumors and their matching tumor-adjacent tissues

**DOI:** 10.1101/2024.06.09.598157

**Authors:** Diana Topolnitska, Camila Lopez Moreno, Alen Paiva, Edward Buchel, Janice Safneck, Afshin Raouf

## Abstract

Therapy resistance and tumor recurrence are major challenges in the clinical management of breast cancer. Current data indicates that the breast tumor microenvironment (TME) and the tumor immune microenvironment (TIME) are important modulators of breast cancer cell response to chemotherapies and the development of therapy resistance. To this end, the ability to recreate the tumor microenvironment in the laboratory using autologous primary cells that make up the breast TME has become an indispensable tool for cancer researchers as it allows the study of tumor immunobiology in the context of therapy resistance. Moreover, the clinical relevance of data obtained from single cell transcriptomics and proteomics platforms would be greatly improved if primary autologous tumor cells were used. In this article, we report a robust and efficient workflow to obtain autologous cancer cells, cancer-associated fibroblasts, and tumor-infiltrating immune cells from primary human breast cancer tumors obtained from mastectomy procedures. As well, we show that this protocol can be used to obtain normal-like epithelial cells, fibroblasts, and immune cells from the matching tumor-adjacent breast tissue samples. Also, a robust methodology to expand each of these primary cell types *in vitro* is presented that allows the maintenance of the primary tumor cell phenotype. The availability of a large number of autologous primary human breast tumor cells and their matching tumor-adjacent tissues will facilitate the study of differential and cancer cell-specific gene expression patterns that will further our understanding of how the TME and TIME influence therapy resistance in the breast tumor context.

## Introduction

Therapy resistance poses a major clinical challenge in the management of estrogen receptor over-expressing (ER^+^) breast cancer tumors as it often leads to tumor recurrence and metastasis^1^. The mechanisms involved in ER^+^ tumor resistance to different endocrine therapies (e.g., fulvestrant and tamoxifen) are multifactorial. In addition to cancer cell-autonomous mechanisms that result in diminishing reliance of breast cancer cells on survival and proliferation signaling pathways that are targeted by cancer therapeutics, therapy resistance in breast tumors is further complicated by the rapidly emerging role of the tumor microenvironment (TME) and the tumor immune microenvironment (TIME)^2,3^. In both cases, paracrine signaling and cell-cell contact interactions between the different cell types that make up the tumor milieu have proven to play essential roles in breast tumor growth, sensitivity to therapeutic agents, and the acquisition of therapy resistance^4-7^. In addition to breast cancer cells, the TME consists of a variety of non-cancerous cells including fibroblasts and their activated derivatives (cancer-associated fibroblasts or CAF), adipocytes, and subsets of immune cells such as the tumor-infiltrating T lymphocytes (TIL)^2,5^. These cell types, together, create a complex and variable milieu that is likely unique to each patient, and plays an essential role in regulating the initial tumor response to therapies and the development of therapy resistance^8,9^.

To date, stable ER^+^ breast cancer cell lines have been extensively used in the study of CAF interactions with cancer cells and the development of endocrine therapy resistance^10,11^. Such studies have revealed valuable insights into supplementary growth survival signals that can be employed by the ER^+^ cancer cells to evade endocrine therapies. However, there are significant drawbacks to the use of these cells in experimentation. For instance, their correlation to clinical breast cancer samples remains poor, and due to significant *in vitro* passaging, they have lost the phenotypic and genotypic heterogeneity of the original tumor sample^12^. This is particularly important with respect to therapy resistance of ER^+^ breast cancer cells, where subsets of cancer cells in the primary tumor may be resistant to therapies but are lost during subsequent passaging *in vitro*^*13*,*14*^. Also, many of the breast cancer cell lines that are used to study endocrine therapy resistance (e.g., MCF7 and T47D) are established from cancer cells that were isolated from pleural effusion samples of lung metastasis of the primary breast tumors^12,13^. Also, these cell lines cannot be used to examine the role of immune cells in the development of therapy resistance as immune cells autologous to these cells are not available. Therefore, the long-established immortalized breast cancer cell lines are not the frank representation of the behaviour of primary cancer cells.

The use of patient-derived breast cancer cells from resected tumors in the study of the TME is steadily increasing. Such assays require complex processes to handle the resected tumor and involve many intricate steps to maximize cancer cell viability and yield. The breast tumor cells can be grown in different 3-dimensional (3D) cultures such as Matrigel^®^ matrix^12,15^. They can also be grown as xenografts in immunodeficient mice^15^. Lastly, primary breast cancer cells can be grown in monolayer 2D cultures^16^. Overall, these culture techniques have significant limitations associated with them. Patient-derived xenografts are expensive to maintain as they require to be passaged *in vivo* within immunodeficient mice^12,15^. Further, the tumor stroma is eventually replaced with murine stromal cells and extracellular matrix^15,17^. Conversely, although 3D cultures can be initiated with freshly resected and dissociated breast tumors, they are limited in the number of times they can be passaged, and the presence of TIL in these cultures is variable at best^15,18^. Lastly, primary breast cancer cells can be expanded in 2D cultures; however, many of the protocols that are currently available are cumbersome and are focused on generating breast cancer cell cultures only and exclude the autologous CAFs and TILs which are important components of the breast TME^19-21^.

Although these culture systems have resulted in exciting advances in understanding the behaviour of cancer cells in the context of their TME, the contribution and study of TIL and other immune cells in the context of their TIME remains challenging^15^. Such important studies require the availability of autologously collected breast cancer cells, CAF, and immune cells, which has proven to be difficult to achieve. Access to such autologous cells will allow for the recreation of the TME as well as some aspects of the TIME *in vitro*. Such autologous cells also provide a framework to study why the current immunotherapies fail to provide complete pathological response against breast tumors. In this article, we describe an optimized and robust protocol to obtain autologous cancer cells, CAF, and immune cells from high-grade ER^+^ breast tumors and their matched tumor-adjacent breast tissue samples obtained from breast-conserving mastectomy procedures, along with an efficient methology to expand these cells *in vitro*.

## Materials and Reagents

### Tissue sample transport to research laboratory from the pathology laboratory

1. Specimen container (Falcon, cat. #354014) with specimen container lid (Falcon, cat. #354017)

### Tissue transport medium

1. Dulbecco’s Modified Eagle Medium (DMEM) with 4500 mg/L Glucose, L-Glutamine, 25 mM HEPES (Gibco™, catalog #12430-054)
2. Heat-inactivated, Fetal Bovine Serum (FBS) (Thermo Fisher Scientific, cat. #12484-028), at 5% of final volume
3. Insulin from bovine pancreas (Millipore Sigma, cat. #I6634), at 10 µg/mL
4. Antibiotic and antifungal solution: prepared by adding 50% (vol/vol) solution of penicillin and streptomycin mix (50X, Gibco™, cat. #15140-122), G418 disulfate salt (50X, Millipore Sigma, cat. #A1720), and Amphotericin B solution (50X, Millipore Sigma, cat. #A2942). This stock solution was stored at -20°C and used at 1X the stock concentration.

### Initial tissue sample processing and dissociation

1. Tissue dissociation shaker flasks, 250 mL (VWR, cat. #71000-426)
2. Stainless-steel surgical blade, No. 22 (Fisher Scientific, cat. # 08-918-5C)
3. Surgical forceps (VWR, cat. #10806-174)
4. Glass Petri dish (VWR, cat. #75845-542)
5. MACS^®^ SmartStrainers, 70 μm (Miltenyi, cat. #130-110-916)
6. Disposable serological pipettes, 5 mL (VWR, cat. #76320-288)
7. Disposable serological pipettes, 10 mL (VWR, cat. #75816-100)
8. Disposable serological pipettes, 25 mL (VWR, cat. #75816-090)
9. Red blood cell lysing buffer, Hybri-Max™ (Millipore Sigma, cat. # R7757)
10. Sharps container, flip-up lid (VWR, cat. #19001-010)

### Tissue dissociation medium (to be prepared immediately prior to use)

1. DMEM plus Nutrient Mixture F-12 (DMEM / F-12) at 1:1 ratio (Gibco™, cat. #11330-032)
2. Bovine Serum Albumin (BSA), lyophilized powder (Millipore Sigma, cat. #A9418), at 2% weight/volume
3. Collagenase from *Clostridium histolyticum* (Millipore Sigma, cat. #C9891), at 300 units/mL
4. Hyaluronidase from bovine testes, Type I (Millipore Sigma, cat. #H3506), at 100 units/mL
5. Insulin from bovine pancreas (Millipore Sigma, cat. #I6634), at 5 μg/mL
6. Hydrocortisone 21-hemisuccinate sodium salt (Millipore Sigma, cat. #H2270), at 0.5 μg/mL
7. Penicillin and streptomycin mix (Gibco™, cat. #15140-122), at 1X the stock concentration

### Cell and/or organoid-like structure freezing medium

1. Cryogenic vials (Corning, cat. #430659)
2. DMEM / F-12 at 1:1 ration (Gibco™, cat. #11330-032)
3. Fetal Bovine Serum (FBS), Heat-Inactivated (Thermo Fisher Scientific, cat. #12484-028), at 46.5% of final volume
4. Dimethyl sulfoxide (DMSO, Fisher Scientific, cat. #BP231-1), at 7% of final volume
5. Nalgene^®^ Mr. Frosty^®^ Cryo 1°C freezing container (Thermo Scientific, cat. #5100-0001)

### Tissue digestion into a single cell suspension

1. Trypsin-EDTA, 0.05% (Gibco™, cat. #25300-062)
2. Dispase in HBSS, 5 units/mL (STEMCELL Technologies, cat. #07913)
3. Deoxyribonuclease (DNase) I from bovine pancreas (Millipore Sigma, DN25), at a final concentration of 100 μg/mL

### Immunomagnetic cell separation

1. OctoMACS™ Separator (Miltenyi, cat. #130-042-109) and MACS^®^ MultiStand (Miltenyi, cat. #130-042-303)
2. MS Columns (Miltenyi, cat. #130-042-201)
3. FcR blocking reagent, human (Miltenyi, cat. #130-059-901)
4. Anti-biotin microbeads (Miltenyi, cat. #130-090-485)
5. Biotin anti-human CD45 monoclonal antibody, clone HI30 (Biolegend, cat. #304004)
6. Biotin anti-human CD31 (PECAM-1) monoclonal antibody, clone WM-59 (Invitrogen, cat. #13-0319-82)

### Collagen solution for coating of tissue culture plates

1. Phosphate Buffered Saline (PBS), pH 7.4 (Gibco™, cat. #10010-023)
2. CellAdhere™ Type I Bovine Collagen Solution, 6 mg/mL (STEMCELL Technologies; cat. #07001), at a final concentration of 0.1 mg/mL

### Breast cancer cell (BCC) expansion medium (concentrations are reported as final concentrations)

1. DMEM / F-12 at 1:1 ratio (Gibco™, cat. #11330-032)
2. FBS (Thermo Fisher Scientific, cat. #12484-028), at 2% of final volume
3. BSA solution, 35% (Millipore Sigma, cat. #A7979), at a final concentration of 1%
4. Human Epidermal Growth Factor (hEGF, Millipore Sigma, cat. #E9644), at 10 ng/mL
5. Insulin from bovine pancreas (Millipore Sigma, cat. #I6634), at 1 μg/mL
6. Hydrocortisone 21-hemisuccinate sodium salt (Millipore Sigma, cat. #H2270), at 0.5 μg/mL
7. (-)-Isoproterenol hydrochloride (Millipore Sigma, cat. #I6504), at 10 μM
8. SB431542 (Hydrate), Transforming Growth Factor Beta (TGF-β) pathway inhibitor (STEMCELL Technologies, cat. # 72234), at 10 μM
9. Y-27632 (Dichloride), RHO/ROCK pathway inhibitor (STEMCELL Technologies, cat. #72304), at 10 μM

### Growth medium for the *in vitro* expansion of CAF or the tumor-adjacent tissue fibroblasts

1. DMEM / F-12 at 1:1 ratio (Gibco™, cat. #11330-032)
2. FBS (Thermo Fisher Scientific, cat. #12484-028), at 10% of final volume

### Expansion of tumor-infiltrating and tumor-adjacent tissue T cells

1. ImmunoCult™-XF T cell expansion medium (STEMCELL Technologies, cat. #10981)
2. Human Recombinant IL-2 (CHO-expressed) (STEMCELL Technologies, cat. #78036), at a final concentration of 10 ng/mL
3. ImmunoCult™ Human CD3/CD28/CD2 T cell activator (STEMCELL Technologies, cat. #10970), at 25 μL/mL of cells

### Indirect immunofluorescent staining reagents

1. 1:1 Acetone (CFS Chemicals, cat. #CFS-104111):Methanol (VWR, cat. #BDH1135-4LP)
2. BSA, Lyophilized Powder (Millipore Sigma, cat. #A9418), at 2% or 5% volume/volume, step-dependent
3. TWEEN^®^ 20 (Millipore Sigma, cat. #P5927)
4. Anti-human alpha-smooth muscle Actin rabbit antibody (abcam, cat. #ab5694)
5. Anti-human vimentin rabbit antibody (Cell Signaling Technology, cat. #3932S)
6. Anti-human S100A4 rabbit antibody (abcam, cat. #ab27957)
7. PE anti-rabbit IgG antibody (Biolegend, cat. #406421)
8. 4′,6-Diamidino-2-phenylindole dihydrochloride (DAPI) (Millipore Sigma, cat. #D8417)

### Reagents for RNA isolation

1. Ethyl alcohol anhydrous, USP (Greenfield Global, cat. #P006EAAN)
2. 2-Mercaptoethanol (Millipore Sigma, cat. #M3148)
3. RNeasy Plus Mini Kit (Qiagen, cat. #74134)
4. Nuclease-Free Water (Promega, cat. #P1193)

### Reagents for cDNA synthesis

1. iScript™ gDNA Clear cDNA Synthesis Kit (Bio-Rad, cat. #1725035)
2. Nuclease-Free Water (Promega, cat. #P1193)

### Polymerase Chain Reaction (PCR) materials and reagents

1. HardShell^®^ 96-well PCR plates (Bio-Rad, cat. #HSP9601)
2. Microseal ‘B’ PCR plate sealing film (Bio-Rad, cat. #MSB1001)
3. TaqMan™ Gene Expression Master Mix (Applied Biosystems, cat. #4369016)
4. *GAPDH* (Glyceraldehyde 3-phsphate dehydrogenase) TaqMan™ probe, Hs02758991_g1 (Life Technologies Inc., cat. #4453320)
5. *ESR1* (Estrogen Receptor alpha) TaqMan™ probe, Hs01046818_m1 (Life Technologies Inc., cat. #4453320)
6. *PGR* (Progesterone Receptor) TaqMan™ probe, Hs01556702_m1 (Life Technologies Inc., cat. #4453320)

### Additional solutions and reagents

1. Ethyl alcohol anhydrous, USP (Greenfield Global, cat. #P016EAAN), diluted to 70%
2. Hanks’ Balanced Salt Solution (HBSS) (Gibco™, cat. #14025-092), supplemented with 2% Fetal Bovine Serum (FBS), Heat-Inactivated (Thermo Fisher Scientific, cat. #12484-028)
3. FITC anti-human CD326 (EpCAM) monoclonal antibody, clone VU-1D9 (STEMCELL Technologies, cat. #60136FI)
4. PE anti-human CD340 (erbB2, HER2) monoclonal antibody, clone 24D2 (Biolegend, cat. #324406)
5. Propidium iodide (PI) solution (Millipore Sigma, cat. #P4864)
6. BD Lyoplate™ Human Cell Surface Marker Screening Panel (BD Biosciences, cat. #560747)

### Additional equipment and materials

1. 1300 Series A2 Class II Biological Safety Cabinet (Thermo Scientific, cat. #1375)
2. Forma™ Steri-Cycle™ CO2 incubator (Thermo Scientific, cat. #370)
3. Heidolph Unimax 1010 shaking incubator (Heidolph Instruments, cat. # 543-12310-04-5) with heating module (Heidolph Instruments, cat. #549-90020-00-3)
4. Heated water bath, at 37°C (VWR, cat. #97025-031)
5. Sorvall™ ST 16R centrifuge (Thermo Scientific, cat. #75004380)
6. Sorvall™ Legend™ Micro 21R microcentrifuge (Thermo Scientific, cat. #75002447)
7. CFX Connect Real-Time PCR Detection System with optical reaction module and thermal cycler (Bio-Rad, cat. #1855201)
8. DNA Engine thermal cycler (Bio-Rad, cat. #ALS-1296)
9. NanoDrop™ 2000C spectrophotometer (Thermo Scientific, cat. #ND-2000C)
10. Vortex-Genie 2 (Scientific Industries Inc., cat. #SI0236)
11. Guava easyCyte™ 8HT flow cytometer (EMD Millipore, cat. #0500-4008)
12. EVOS™ M5000 Imaging System (Invitrogen, cat. #AMF5000), and EVOS™ XL Core Configured Cell Imager (Invitrogen, cat. #AMEX1100)
13. Automatic-Sarpette^®^ serological pipetting aid (Sarstedt, cat. #90.189.203)
14. Eppendorf Research^®^ 100 – 1000 μL adjustable-volume pipette (Eppendorf, cat. #022472101), 20 – 200 μL pipette (Eppendorf, cat. #022472054), 2 – 20 μL pipette (Eppendorf, cat. #022471953), and Finnpipette F1 0.2 – 2 μL pipette (Thermo Scientific, cat. #4641010)
15. 1000 μL Redi-Tip™ pipette tips (Fisher Scientific, cat. #02-681-163), 200 μL pipette tips (Axygen, cat. #301-02-301), and 10 μL pipette tips (Axygen, cat. #301-03-051)
16. 1250 μL filtered pipette tips (FroggaBio, cat. #FT1000), 200 μL filtered pipette tips (FroggaBio, cat. #FT200), 10 μL Finntip™ filtered pipette tips (Thermo Scientific, cat. #94052100), and 10 μL uTIP™ filtered pipette tips (VWR, cat. #89136-572)
17. 1.7 mL microtubes (Axygen, cat. #MCT-175-A) and 0.2 mL microtubes (Axygen, cat. #PCR-02-A)
18. Round-bottom polystyrene test tubes, 5 mL (Falcon, cat. #352054)

11 15 mL conical centrifuge tubes (Falcon, cat. #352097), and 50 mL conical centrifuge tubes (Thermo Scientific, cat. #339652)
12 96-well clear polystyrene microplates (Corning, cat. #3797)
19 6-well polystyrene microplates (Falcon, cat. #353046), and 10 cm TC-treated culture dishes (Corning, cat. #430167)
20 Nitrile gloves, small (Kimberly-Clark Professional™, cat. #50706), medium (Kimberly-Clark Professional™, cat. #50707), and large (Kimberly-Clark Professional™, cat. #50708)
21 Tyvek™ sleeves (DuPont™, cat. #TY500SWH00020000)

## Detailed Protocol

### Tissue procurement

1 The breast tumor and the matching tumor-adjacent tissue (TAT) were collected from breast cancer patients with tumors over 2 centimeters in size that were undergoing breast conserving mastectomy surgeries in accordance with approved ethics protocols of the University of Manitoba (HS25018). The TAT samples were collected from tissue located over 3 centimeters away from the primary tumor boundary. The tumor samples were collected based on the discretion of a breast cancer pathologist. For the best results, specimen containers filled with the transport medium were made available in the pathology laboratory in advance of the surgery. The tumor and the matching TAT samples were placed in the transport medium immediately after pathological examination. In the case of mastectomy surgeries that were followed by breast reconstructive procedures, unused fat samples from abdominal flaps autologous to the breast tumor and TAT were collected and processed as described^2,3^. These fat samples are a valuable source of autologous tissue-resident immune cells, adipose-derived stem cells, and normal (non-activated) fibroblasts. For the purposes of this study, all breast tumors were ER^+^ and locally invasive (**Table 1**).

**Table 1.** Clinical sample summary and phenotype of the *in vitro* expanded primary cells. 8 ER^+^ breast tumor samples were obtained and dissociated. The BCC and CAF were separately expanded *in vitro* (see text for detailed protocol). The primary tumor diagnosis was obtained from the pathology reports. ERα (encoded by the *ESR1* transcript) and PR (encoded by the *PGR* transcript) expression of the *in vitro* expanded BCC was obtained by RT-qPCR. HER2 protein expression was examined by flow cytometry. LN, lymph node; ER, estrogen receptor; PR, progesterone receptor; EpCAM, epithelial cell adhesion molecule; CAF, cancer-associate fibroblasts; BCC, breast cancer cells; TAT, tumor-adjacent tissue; F, fibroblasts; BEC, breast epithelial cells, Y, yes; N, no; N/A, not available.

| Sample ID | Primary tumor diagnosis | CAF obtained | BCC obtained | Tumor CD45/CD31 cells obtained | BCC characterization | Frequency of EpCAM+ BCC | TAT-F obtained | TAT-BEC obtained | TAT CD45/CD31 cells obtained | TAT-BEC characterization | Frequency of EpCAM+ TAT-BEC |
| --- | --- | --- | --- | --- | --- | --- | --- | --- | --- | --- | --- |
| S1 | ER+ PR+ HER2-, LN-negative | N | Y | Y | ER+ PR+ HER2- | 82.8 | N | N | Y | N/A | N/A |
| S2 | ER+ PR+ HER2-, LN-negative | N | Y | Y | ER+ PR+ HER2- | 15.4 | N | N | Y | N/A | N/A |
| S3 | ER+ PR+ HER2-, N2, LN-negative | N | Y | Y | ER+ PR- HER2- | 33.7 | N | N | Y | N/A | N/A |
| S4 | ER+ PR+ HER2-, LN-negative | Y | Y | Y | ER+ PR+ HER2- | 7.02 | N | Y | Y | ER+ PR- HER2- | 90.8 |
| S5 | ER+ PR+ HER2+, LN-positive | N | Y | Y | ER+ PR- HER2- | 94.6 | Y | Y | Y | ER+ PR- HER2- | 93.6 |
| S6 | ER+ PR+ HER2-, LN-negative | Y | Y | Y | ER+ PR+ HER2- | 96.0 | Y | Y | Y | ER+ PR+ HER2- | 97.7 |
| S7 | ER+ PR+ HER2-, LN-positive | Y | N | Y | N/A | N/A | N | Y | Y | ER+ PR- HER2- | 90.8 |
| S8 | ER+ PR+ HER2-, LN-N/A | Y | Y | Y | ER+ PR- HER2- | 98.5 | Y | N | Y | N | N/A |

*<u>Note A</u>*: *Sample cups containing transport medium can stay at room temperature for up to 3 hours*.

*<u>Note B</u>*: *In the case of mastectomies that are followed by reconstructive surgeries, the unused fat graft can be collected in a specimen cup containing the transport medium*.

*<u>Note C</u>*: *It is best to prepare fresh transport medium prior to each sample collection*.

2 The specimen containers were transported to the research laboratory and immediately placed in a 4°C fridge until further processing.

*<u>Note A</u>*: *Tissue samples can be kept in the transport medium for up to 2 days (e*.*g*., *over the weekend) as long as they remain completely submerged in the growth medium. However, for the best results, it is highly recommended that samples be processed on the day of collection*.

*<u>Note B</u>*: *If the samples are to be processed immediately upon collection, it is important to leave the tissues in the transport medium for at least 1 hour to limit any bacterial and/or fungal contamination*.

### Tissue processing and dissociation

1 The collected samples were processed in a biosafety cabinet (BSC). A sterile environment was created by following the manufacturer’s protocol. In addition, the work surface inside the BSC unit was wiped clean with 70% ethanol. All procedures from here on were performed in a sterile environment, using sterile reagents and sterile tissue culture techniques to limit contamination of the samples.
2 The tumor or the adjacent tissue samples were removed from the specimen cups and placed in separate glass Petri dishes where they were minced with scalpels.

*<u>Note</u>*: *If sample size allowed, a small portion of each tissue was extracted prior to being minced, flash frozen, and kept in a -80°C freezer for histological and other analyses*.

3 The minced tissue samples were placed in dissociation medium in separate flasks and shaken using an orbital shaker at 100-110 revolutions per minute (RPM) at 37°C for 16 hours.

*<u>Note A</u>*: *For best results, the RPM and temperature settings should be optimized for each orbital shaker to avoid over/under-dissociation of the tissue samples. These optimization steps are required to minimize cell loss and maintain optimum enzyme activity*.

*<u>Note B</u>*: *The dissociation time (16 hours) should also be optimized for each instrument. Further, using an orbital shaker that provides consistent temperature throughout the tissue dissociation step is required to prevent over-digestion of the samples and maintaining cell viability*.

*<u>Note C</u>*: *Small organoid-like structures, or tissue clumps, may still be visible following dissociation due to the fibrotic nature of some breast tumor and the tumor-adjacent tissue samples*.

4 The organoid-like structures and released cells were pelleted via centrifugation at 1200 RPM for 5 minutes at room temperature.

*<u>Note</u>*: *If stringy, connective tissue is observed after overnight dissociation, do not extend dissociation time in the orbital shaker. Using a 1 mL pipette, attempt to breakdown the stringy bits and ultimately remove them before proceeding to the next step*.

5 The pellets containing the organoid-like structures and released cells were washed in Hanks’ Balanced Salt Solution (HBSS) supplemented with 2% fetal bovine serum (FBS), 2%HBSS, and pelleted again via centrifugation at 1200 RPM.
6 At this point, the pelleted organoid-like structures and released cells can be cryopreserved in the freezing medium, transferred to a Mr. Frosty™ container and placed in a -80°C freezer. After 24 hours, the frozen samples can be placed in the vapor phase of a liquid nitrogen tank for long-term storage.

### Creating single cell suspensions from dissociated breast tumor and tumor-adjacent tissue samples

1 Organoid-like structures and released cells (either from freshly dissociated samples or previously frozen pellets) were placed in a 15 mL centrifuge tube and dissociated with 1 mL of warm trypsin for 7 minutes in a 37°C water bath. Then after, trypsin was deactivated with 1 mL of 2%HBSS prior to being pelleted via centrifugation at 1200 RPM at room temperature.

*<u>Note A</u>*: *When thawing frozen samples a parafilm strip should be wrapped around the lid of the cryoviasl to protect against the spread of plastic pieces in the unlikely event that the vials were to explode due to the sudden temperature change. The vials were placed in a 37°C water bath until a small ice pellet is still visible. The vials where the transferred to a BSC unit and their contents was transferred to a 15 mL centrifuge tube containing 5 mL of* 2% HBSS. *Cells were then pelleted via centrifugation at room temperature at 1200 RPM. To ensure cell viability is maintained, extra care must be taken to thaw the cells as efficiently as possible as the DMSO component of the freezing medium is toxic to the cells*.

*<u>Note B</u>*: *In instances where tissue sample and subsequent cell pellet size is large, up to 2 mL of trypsin can be added to aid in dissociation*.

2 The pellet was further dissociated with 1 mL of warm (37°C) dispase solution for 7 minutes in a 37°C water bath. An equal volume of 2%HBSS was then added to the cell suspension prior to being pelleted via centrifugation at 1200 RPM at room temperature.

*<u>Note</u>*: *It is important not to leave the dispase solution in the warm water bath for more than 10 minute as this could result in enzyme deactivation*.

3 The cell pellet was then re-suspended in 10 mL of ice-cold 2%HBSS solution and passed through a 70 µm mesh placed on top of a 50 mL centrifuge tube. The filter mesh was washed using 5 mL 2%HBSS and the cells were pelleted via centrifugation at 1200 RPM at room temperature. The cell pellet was resuspended in 1 mL of ice-cold 2%HBSS.

*<u>Note</u>*: *To ensure a single cell suspension has been generated, the cell pellet should be pipetted repeatedly in the* 2%HBSS *with a 200 µL pipette tip*.

4 10 µL of the cell suspension was placed on a glass slide and observed under an inverted microscope to ensure that <5% of cells are in aggregates (>3 cells attached together). If

>5% of the cell suspension contained aggregates, steps 1-3 were repeated.

*<u>Note</u>*: *At this point, the cells can be used to generate organoids in 3-dimensional cultures as described*^*22-24*^*or other experiments, as needed. Alternatively, the cells can be pelleted and cryogenically stored using the procedure described*.

5 If the cell pellet obtained upon dissociation is visibly red, the cells can be treated with red blood cell lysis buffer as per the manufacturer’s protocol.

### Immunomagnetic separation of the hematopoietic and endothelial cell fraction

1 The single cell suspension from the previous step was depleted of hematopoietic (CD45-expressing) and vascular endothelial (CD31-expressing) cell lineages using immunomagnetic cell separation. To start, cells were resuspended in 2%HBSS to which the FcR blocking reagent and biotin conjugated anti-CD45 and anti-CD31 antibodies were added as per manufacturer’s protocol. This mixture was allowed to incubate at room temperature for 15 minutes. Next, the cells were washed twice and resuspended in 2%HBSS.

*<u>Note A</u>*: *Antibody dilutions should be optimized for each application*.

*<u>Note B</u>*: *During wash steps, the cells should be centrifuged at 1200 RPM for 5 minutes at room temperature, as before*.

2 Anti-biotin microbeads were added at the manufacturer-recommended dilution and following a 15-minute incubation at 4°C, the cells were washed and centrifuged at 1200 RPM for 5 minutes at room temperature. Finally, the cells were resuspended in an appropriate volume of 2%HBSS for immunomagnetic separation, following the manufacturer’s recommendation.

*<u>Note</u>*: *For optimal results, it is recommended that the cells be incubated with the microbeads in a 4°C refrigerator. Avoid incubation on ice*.

3 MS columns were prepared and used as recommended by the manufacturer. This procedure separates the tumor cell suspension into the lineage-depleted cells and the CD45^+^/CD31^+^ cell mixture (**Fig. 1A**).

**Figure 1.**
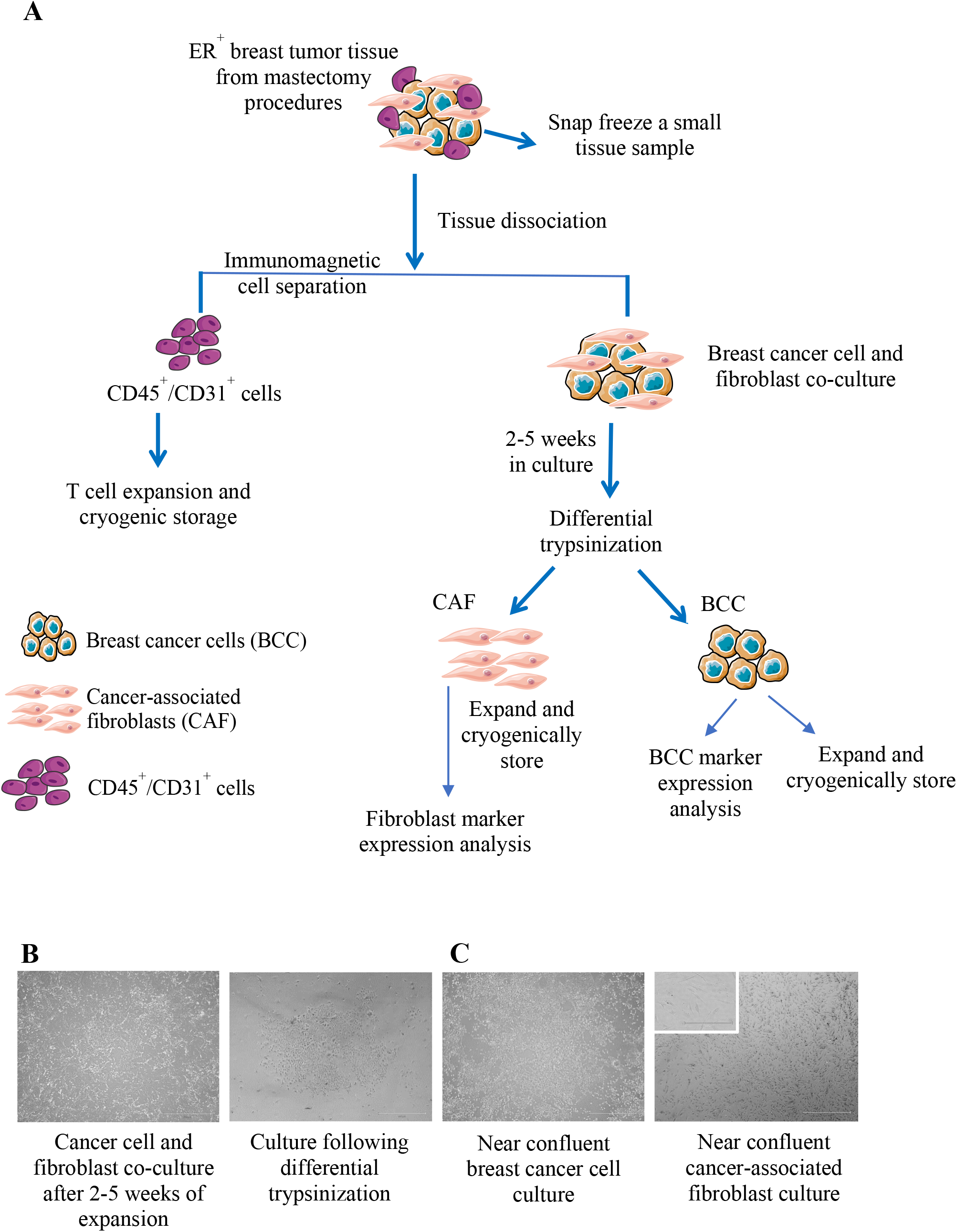
Primary breast cancer cell cultures following differential trypsinization. (**A**) Flow chart depicting breast tumor sample handling and dissociation. ER^+^ breast tumors (**Table 1**) were obtained from the Pathology laboratory, dissociated and placed in collagen-coated tissue culture plates (see text for detailed protocol). After 2 – 5 weeks, the cancer-associated fibroblasts (CAF) were removed via differential trypsinization and grown in a separate collagen-coated plate using DMEM / F12 growth medium supplemented with 10% FBS. The breast cancer cells (BCC) were allowed to continue to expand in their original tissue culture plates. (**B**) Representative images illustrate BCC–CAF co-cultures and the BCC colonies that follow differential trypsinization. (**C**) BCC and CAF were allowed to grow in culture until 70% confluent using their specialized growth medium, as described. Representative images are shown. The scale bars represent 300 µm and 600 µm.

*<u>Note A</u>*: *The magnetically labeled cells are flushed by firmly pushing a plunger into the column. To ensure that all cells are released from the column, this step should be done twice*.

*<u>Note B</u>*: *To increase the purity of the lineage negative fraction, the cells can be passed over a second column*.

*<u>Note C</u>*: *Immunomagnetic cell separation results in significant cell losses from the positively selected cells. It is highly advisable that several dry run experiments be*

*conducted to ensure proficiency in performing this step and to adhere closely to the manufacturer’s protocol*.

*<u>Note D</u>*: *Fluorescent activated cell sorting (FACS) can also be used to remove CD45/CD31-expressing cells from the tumor cell suspension. However, most tumor samples used to develop this protocol did not yield sufficient cell numbers to perform efficient cell separation via FACS*.

4 The CD45^+^/CD31^+^ cells obtained in the previous step were cryogenically preserved at this point (up to 1×10^6^ cells/cryovial), as described. These CD45^+^/CD31^+^ cells can be used in studies pertaining to the biology of tumor-infiltrating lymphocytes.

*<u>Note A</u>*: *The tumor-infiltrating T cells were expanded from the CD45*^*+*^*/CD31*^*+*^ *cells using IL-2 supplemented ImmunoCult™-XF T cell expansion medium. The T cells were stimulated using the ImmunoCult™ CD3/CD28/CD2 T cell activator. In our experience, the CD31*^*+*^ *cells were eliminated in these cultures as the conditions are not suitable for their survival and proliferation (data not shown).*

*<u>Note B</u>*: *The expanded T cells were cryogenically preserved for future experiments*.

### Obtaining autologous breast cancer cells and cancer-associated fibroblasts from the dissociated tumor samples

1 The lineage-depleted (CD45/CD31-removed) tumor cells were pelleted via centrifugation at 1200 RPM at room temperature and the cell pellet was then resuspended in breast cancer cell (BCC) expansion medium. In the meantime, a 10 cm tissue culture plate was coated with diluted bovine collagen solution for 20 minutes at 37°C in a humidified incubator with 5% CO2. After 20 minutes, the tissue culture plates were washed with 3 – 5 mL of PBS to remove any unpolymerized collagen.

*<u>Note A</u>*: *Enough diluted collagen should be added to each plate to compensate for liquid evaporation during the incubation period. For a 10 cm tissue culture plate, a minimum of 3 mL of collagen solution should be used*.

*<u>Note B</u>*: *Care must be taken not to leave the plates in the incubator for more than 30 minutes to avoid drying out the polymerized collagen*.

*<u>Note C</u>*: *Depending on the concentration of collagen per batch, the dilution factor may need to be adjusted. This step is important, and care must be taken to achieve consistent collagen concentration and plate coverage*.

2 The dissociated tumor cell suspension was added to a collagen-coated plate containing 10 mL of BCC growth medium. The plate was placed in a humidified incubator and left undisturbed until colonies consisting of cancer-associated fibroblasts (CAF) and BCC were observed. The growth medium was changed every 3^rd^ day (**Fig. 1B**).

*<u>Note A</u>*: *Initially, the BCC grew as separate colonies, visible 2-5 weeks after culture initiation. With passing days, CAF became distinctly visible in the space surrounding the BCC colonies*.

*<u>Note B</u>*: *Due to the extended time required to establish these cultures, exceptional tissue culture hygiene practices are recommended (e*.*g*., *using separate BSC units, humidified incubators and tissue culture plastics and reagents are encouraged)*.

*<u>Note C</u>*: *A stock solution of BCC growth medium can be prepared with all components except hEGF, TGF-β and RHO/ROCK pathway inhibitors. This stock solution is stable for up to 14 days. Once hEGF and the inhibitors are added to the medium, it should be used within 7 days*.

3 When the 10 cm plate(s) reached near confluency, CAF were lifted using the differential trypsinization technique^25^ with some modifications. Briefly, 2 mL of warm (37°C) trypsin was added to each plate and allowed to incubate at 37°C for 2 minutes. Then after, the plate was gently taped and rinsed with an equal volume (2 mL) of 2%HBSS (**Fig. 1B**).

*<u>Note</u>*: *In instances where CAF may be resistant to be lifted, the differential trypsinization step should not exceed 3 minutes to avoid dislodging the cancer cells. See other notes to remove any remaining CAF from the BCC cultures*.

4 The dislodged cells were transferred to a centrifuge tube and pelleted at room temperature at 1200 RPM for 5 minutes, suspended in 10 mL of DMEM growth medium supplemented with 10% FBS, and plated in a new 10 cm collagen-coated tissue culture plate.

*<u>Note A</u>*: *At this point, a cell count should be done to ascertain the number of fibroblasts that were recovered from the culture plates. In our experience, up to 3x10*^*5*^ *CAF can be plated into 1 collagen-coated 10 cm plate*.

*<u>Note B</u>*: *In our experience, re-plating the lifted fibroblasts on a collagen-coated tissue culture plates improves the efficiency of establishing CAF cultures. See Discussion section for more details*.

5 The CAF cultures should be passaged once they reach 70 – 75% confluency (**Fig. 1C**). The CAF can be lifted with warm trypsin using the previously published techniques^26^.

*<u>Note A</u>*: *It is essential that the CAF cultures do not exceed 75% confluency as this will shorten their expansion potential*.

*<u>Note B</u>*: *At every passage, CAF should be cryogenically frozen for future use. We do not recommend using primary CAF beyond passage 6*.

*<u>Note C</u>*: *Starting with the second passage, the CAF should be characterized with respect to their expression of markers associated with an activated fibroblast phenotype (e*.*g*., *alpha-smooth muscle actin*^*27*^).

*<u>Note D</u>*: *It is essential to monitor the presence of Epithelial Cell Adhesion Molecule expressing (EpCAM*^*+*^*) cells in the CAF cultures as they represent the contaminating breast cancer cells*^*28*^. *Based on our experience, the EpCAM*^*+*^ *cells should be nearly undetectable (****Supp. Fig. 1****) in the CAF cultures by passage 3. However, if >10% of CAF cultures are EpCAM*^*+*^, *differential trypsinization (steps 3-4) should be repeated*.

### *In vitro* expansion of primary human breast cancer cells

1 Following the removal of the CAF, the tissue culture plates containing BCC colonies were washed with 2%HBSS to remove any loose CAF cells and allowed to grow in BCC growth medium until the plates either reached 70 – 80% confluency or the center of the colonies became over-confluent (**Fig. 1C**).

*<u>Note A</u>*: *It is important to monitor these cultures closely as BCC expand. If necrotic centers begin to form, the BCC cultures should be passaged to avoid further cell loss*.

*<u>Note B</u>*: *A small portion of the cells were stained with a fluorescently tagged anti-human EpCAM antibody to obtain an approximate breast cancer (EPCAM*^*+*^*) cell number using a flow cytometer. This is only an approximation because the percentage of EpCAM*^*+*^ *cells in breast cancer cells obtained from different tumors is highly variable (****Table 1****). Because we have focused on ER*^*+*^ *tumors, however, the percentage of EpCAM*^*+*^ *cells were found to be frequently high*.

*<u>Note C</u>*: *At this stage, some BCC cultures were observed to contain fibroblast-like cells. In such cases, the differential trypsinization steps were repeated to improve the purity of the BCC cultures*.

2 During expansion, the growth medium was replaced every 2 – 3 days.

*<u>Note A</u>*: *During the first week of culture, extreme care must be taken not to significantly disturb the BCC when changing the growth medium. We recommend leaving 10% of the spent medium in the plates during media change*.

*<u>Note B</u>*: *Each culture plate should be observed under an inverted microscope to examine cell shape and cell size. Most cells should have a cuboidal shape, which is typical of ER*^*+*^ *BCC. At later passages, nearly all cells should adopt a cuboidal shape (****Fig. 1C***).

### *In vitro* passaging of primary human breast cancer cells

1. BCC cultures were passaged once they reached 70% – 80% confluency.
2. The growth medium was removed from the cultures and the BCC were washed once with warm PBS at 37°C.
3. Warm trypsin (2 mL) was added to cover the 10 cm plate’s surface and the plate was placed in a humidified incubator with 5% CO2 for 5 – 7 minutes at 37°C.
4. The plates were tapped gently to free loosened cells and a pipette with 1 mL tip was used to rinse the bottom of the plates several times to dislodge as many cells as possible. Plates were observed under an inverted microscope to ensure most cells were lifted.

*<u>Note A</u>*: *If the BCC prove difficult to lift from the plates, warm dispase can be used in combination with trypsin at equal volume*.

*<u>Note B</u>*: *In instances where cells are more resistant to lift, the incubation time can be extended up to 12 minutes at 37°C, with occasional tapping and constant monitoring under a microscope. This step, however, should not exceed 12 minutes as cell viability will be significantly decreased*.

5 The trypsinization process was stopped by adding an equal volume (2 mL) of 2%HBSS to the plates. The bottoms of the plates were rinsed using a 1 mL pipette. The plates were observed under an inverted microscope to ensure most cells were released from the plate.
6 The released cells were pelleted via centrifugation at 1200 RPM for 5 minutes at room temperature, and the cell pellet was resuspended in 1 mL 2%HBSS. The cells were counted using a hemocytometer or an automated cell counter.
7 To continue to propagate the BCC, 5×10^5^ cells were plated into a collagen-coated 10 cm tissue culture plate, as described.

*<u>Note A</u>*: *In our experience, the BCC started to grow more rapidly after the first passage. Therefore, it is important to check the cultures every day under a microscope and change their growth medium every 2 – 3 days, as required*.

*<u>Note B</u>*: *Always freeze cells from early passages for future experiments*.

*<u>Note C</u>*: *The BCC should be monitored for expression of fibroblast markers (e*.*g*., *alpha-smooth muscle actin, SMA). After passage 3, no fibroblasts should be detected in the cultures. If fibroblast-like cells (SMA*^*+*^ *cells) are still present in these cultures after the third passage, the cultures can be subjected to differential trypsinization, as described*.

*<u>Note D</u>*: *It is recommended that the BCC be characterized at different passages to ensure that their phenotype resembles that of the original tumor in terms of ER, PR, and HER2 expression (see “Characterization of in vitro expanded breast cancer cells” section)*.

*<u>Note E</u>*: *It is not recommended to use primary BCC beyond passage 7*.

### Obtaining autologous cells from the tumor-adjacent tissue

The matching tumor-adjacent tissue (TAT) to each breast tumor was dissociated using the steps described for dissociating breast tumors. The TAT -epithelial cells, -immune cells and -fibroblast cells were obtained using the methodology outlined above.

*<u>Note</u>*: *In our hands, TAT-fibroblasts also require to be plated on collagen-coated plates. Precautions listed above regarding the care for the cells and cell culture hygiene must be observed*.

### Characterization of *in vitro* expanded breast cancer cells

The methodology described here, has been optimized through an iterative process. We initially plated the CD45/CD31 lineage-depleted tumor cells for 3 – 4 hours in non-collagen coated tissue culture plates, allowing CAF to adhere to the plate. Then after, the non-adherent cells were plated in collagen-coated culture plates with the BCC growth medium, while the adhered CAF were grown in DMEM medium supplemented with 10% FBS. This procedure was based on the differential sedimentation rate of fibroblasts compared to the breast cancer cells^29^. Although this methodology worked well to allow the growth of BCC, it was inefficient in generating CAF cultures. Using this procedure, we generated and expanded primary BCC lines from 4 individual ER^+^ breast tumors, however, only 1 CAF line could be obtained. We, then, modified this procedure to the method detailed above, and were able to successfully expand 3 BCC, 3 CAF, and 4 TIL lines from 4 ER^+^ tumors (**Table 1**). Also, using this modified procedure, we successfully obtained and expanded primary breast epithelial cells (TAT-BEC, 3 lines), immune cells (4 lines) and fibroblasts (TAT-F, 3 lines) from the 4 patient-matched TAT samples (**Table 1**). Although both methods outlined allowed for the generation of primary BCC lines (7 lines established out of 8 samples attempted), only the differential trypsinization technique and the use of collagen-coated culture plates favored the development of primary CAF and TAT-F lines in an efficient manner.

To examine if the original BCC and CAF phenotypes observed in the tumors were maintained following their *in vitro* passaging, we examined the expression of biomarkers specific to each cell type. In the case of fibroblasts, the high expression of vimentin, fibroblast-specific protein 1 (S100A4) and alpha-smooth muscle actin (αSMA) was monitored as an indication of a high frequency of activated phenotype in the CAF (**Fig. 2A**). As expected, the TAT-F contained fewer activated fibroblasts, which was discerned based on their decreased expression of αSMA (**Fig. 2A** and references ^6,22^). The phenotype of the expanded BCC lines was examined at passages 5 – 7 by quantifying their expression of estrogen receptor alpha (ERα, encoded by *ESR1*) and progesterone receptor (PR, encoded by *PGR*) using RT-qPCR. RNA from BCC were made into cDNA and the expression of different biomarkers were obtained using TaqMan^TM^ probes. Human epidermal growth factor receptor 2 (HER2) protein expression was examined by flow cytometry (**Fig. 2B and 2D**). All established primary BCC lines expressed high levels of *ESR1* with variable *PGR* transcript levels (**Fig. 2B**). This data was further corroborated by the variable expression of EpCAM in these lines (**Table 1**). All BCC samples showed low expression of HER2 protein (**Fig. 2D**). Interestingly, the tumor sample S5 was found to be HER2^+^ based on the pathology report, but the *in vitro* expanded S5-BCC were found to have low expression of HER2 protein (**Fig. 2D**). The patient-matched TAT was also obtained for 5 of the samples studied (S4 – S8). All samples apart from one, S8, yielded TAT breast epithelial cells (TAT-BEC) that could be expanded in culture (**Table 1**). We further characterized these TAT-BEC and found that they have high expression of ERα and PR but are negative for the expression of the HER2 protein (**Fig. 2C and 2D**). Interestingly, unlike the BCC, expanded TAT-BEC consistently showed high expression of EpCAM (**Table 1**).

**Figure 2.**
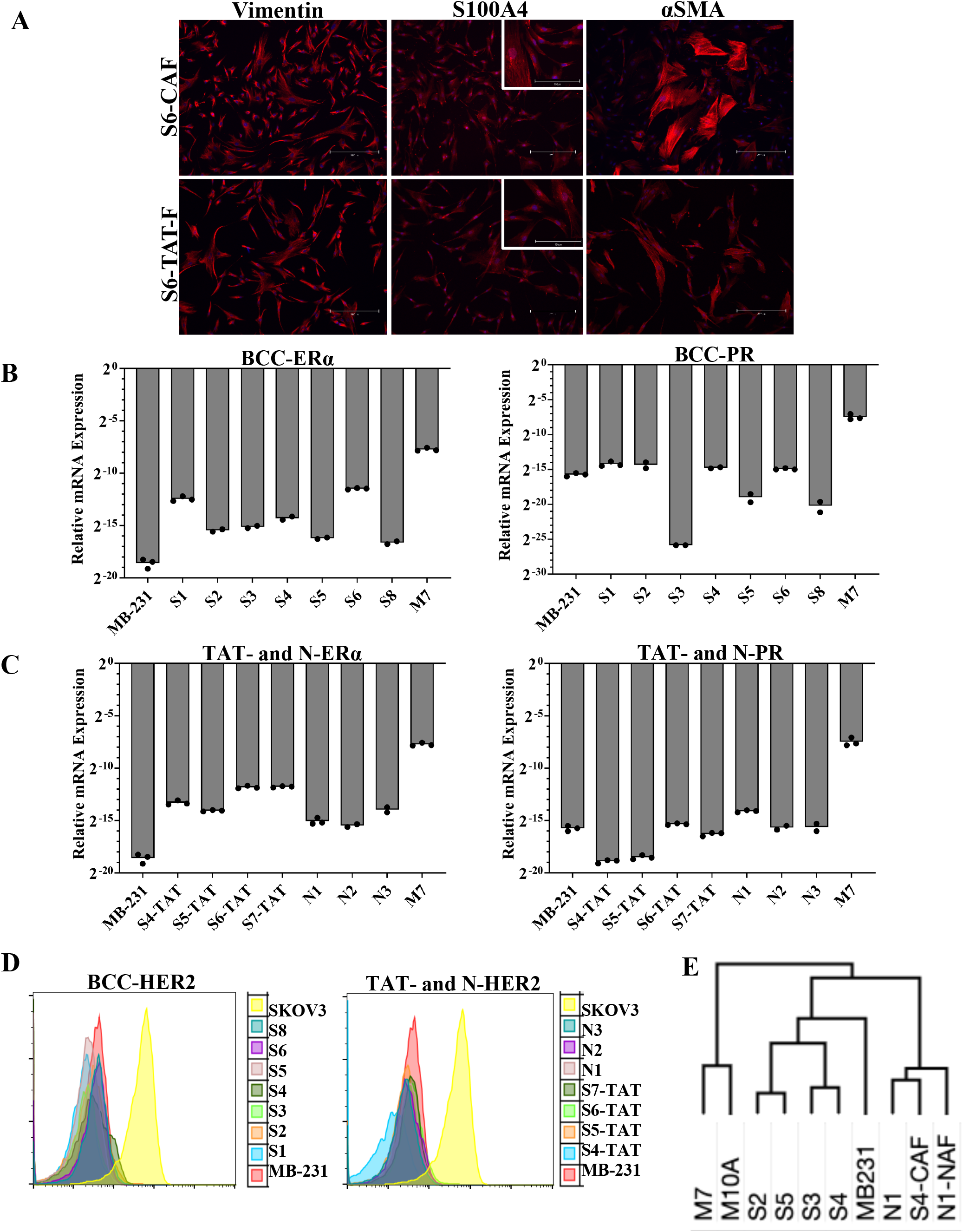
Characterization of primary breast cancer cells and matching fibroblasts. Breast tumors were dissociated and the breast cancer cells (BCC) and cancer-associated fibroblasts (CAF) were separately expanded *in vitro* for approximately 5 passages. From the matching tumor-adjacent tissue (TAT), fibroblasts (TAT-F) and breast epithelial cells (TAT-BEP) were also separately expanded *in vitro*. (**A**) Protein expression of vimentin, fibroblast-specific protein 1 (FSP-1, also called S100A4), and alpha-smooth muscle actin (αSMA) were used to determine the activated phenotype of the CAF and TAT-F using immunofluorescence. The scale bars represent 150 µm and 300 µm. (**B**) The expression of ERα (encoded by *ESR1*) and PR (encoded by *PGR*) in BCC, (**C**) TAT-BEP (TAT), BEP from normal breast tissue (N), MCF7 (M7) cells, and MDA-MB-231 (MB-231) cells was determined using RT-qPCR. Transcript expression was normalized to *GAPDH* transcript levels in all samples. (**D**) The expression of the HER2 receptor was determined using flow cytometry. Here, SKOV3 and MDA-MB-231 cells were used as positive and negative controls, respectively. (**E**) The expression of 243 cell surface markers was examined using flow cytometry on the surface of *in vitro* expanded BCC, CAF, normal breast epithelial cells (N), and normal-associated fibroblasts (NAF). MCF7 (M7), MCF10A (M10A), and MDA-MB-231 (MB231) cells were also examined for their marker expression. The data was analyzed using FlowJo Version 9.9.5. The frequency of cells positive for the expression of each receptor (**Supplementary Table 2**) was used to generate a hierarchical cluster of the samples. For this purpose, the Pearson correlation was used with complete linkage method (Morpheus, https://software.broadinstitute.org/morpheus).

We had previously developed a growth medium similar to the BCC expansion medium (detailed in the Materials and Reagents section) for the generation of primary normal human breast epithelial cells from reduction mammoplasty samples (**Table 2**)^7^. Therefore, to assess the level of contaminating normal breast epithelial cells in the BCC lines, we examined the expression of 244 cell surface receptors on 5 primary BCC lines and 1 primary normal human breast epithelial cell sample obtained from discarded mammoplasty tissue. These lines were chosen as they produced the high cell numbers (∼14×10^6^ cells) that were required to carry out the cell surface receptor array. For this purpose, we used an array of antibodies raised against human cell surface receptors, which were detected using a secondary antibody conjugated to Alexa Fluor™ 647, based on the manufacturer’s recommendations (BD Biosciences). In this way, we quantified the frequency (percent positivity) of each cell surface receptor (**Supp. Table 1**). We, then, employed hierarchical clustering analysis on the receptor expression data and generated a heatmap to evaluate how closely the BCC lines are related to each other and to the normal breast epithelial cells (**Fig. 2E**). The primary normal epithelial cells (N1) and the two fibroblast lines (S4-CAF and N1-NAF) clustered separately from the primary BCC lines, which, on their own, formed 2 separate clusters. Interestingly, the long-established breast cancer and the normal-like breast epithelial cell lines (MCF7 and MCF-10A, respectively) formed a cluster separate from the primary BCC and the normal breast epithelial cells (**Fig. 2E**). This data indicates that the primary BCC lines generated are of high purity and contain little contaminating fibroblasts or normal epithelial cells.

**Table 2.** Summary of the normal breast tissue samples and the phenotype of the *in vitro* expanded normal epithelial and fibroblast cells. Discarded samples from 3 reduction mammoplasty surgeries were dissociated, and the normal breast epithelial cells (N-BEC) were expanded as previously described ^7^. The phenotypes of the *in vitro* expanded N-BEC were obtained by RT-qPCR (*ESR1* and *PGR* transcript expression) and flow cytometry (HER2 protein expression).

| Sample ID | Pathology Report | N-BEC obtained | N-BEC characterization | Fibroblasts obtained |
| --- | --- | --- | --- | --- |
| N1 | Normal, mammoplasty sample | Y | ER+ PR+ HER2- | Y |
| N2 | Normal, mammoplasty sample | Y | ER+ PR- HER2- | N |
| N3 | Normal, mammoplasty sample | Y | ER+ PR- HER2- | N |

## Discussion

Here, we present a robust yet simple workflow for procurement of breast tumor samples, tumor dissociation, and the cell culture conditions required for the expansion of the different cell types that make up the breast tumor microenvironment. In this study, the tumor samples were all ER^+^ luminal breast tumors as they make up over 75% of the diagnosed cases of breast cancer. The growth medium we used here consists of TGF-β and RHO/ROCK pathway inhibitors to minimize the epithelial to mesenchymal transition (EMT) of the ER^+^ breast cancer cells in these cultures. This growth medium also contains a small percentage (2% of final volume) of FBS, which is the likely source of estrogen needed for ER^+^ breast cancer cell survival and proliferation. It is interesting that the best results were obtained when BCC and CAF were cultured together during their early *in vitro* growth phase (P0 – P1). This environment likely provides essential survival signals for the cancer cells through cell–cell contact and paracrine communications, which have been shown to play important roles in the breast tumor niche. Using the outlined methodology, the BCC lines generated were shown to contain minimal contaminating CAF or normal breast epithelial cells, which makes these BCC suitable for the study of ER^+^ tumor biology, their intrinsic resistance to therapies, and the development of therapy resistance. It is interesting that the long-established ER^+^ breast cancer cell line MCF7 shared very little similarity with the primary ER^+^ BCC lines. However, this is not entirely surprising as MCF7 cells were derived from the pleural effusion of an individual with metastatic ER^+^ breast cancer^30^ and the high passage number of MCF7 cells may have caused a drift in their gene expression profiles^30^. Since the protocol outline here allows for generation of sufficient cell numbers needed at low passages (i.e., passage 4 – 5), these cells may phenocopy the primary breast cancer cell behavior *in vivo*. The protocol outlined here is not perfect however, as not all BCC lines were capable of generating the large number of cells needed for cell surface receptor array profiling experiments. Moreover, S5-BCC do not phenocopy the tumor that they originate from in terms of their surface HER2 expression levels. This could be due to the rapid expansion of the HER2– cell clones as compared to the HER2-expressing clones. Also, although the primary tumor was deemed to be HER2^+^ based on pathology report, the expression of HER2 in the original tumor may have been low. In any case, this protocol may not be suitable for expansion of BCC from primary ER^+^HER2^+^ or HER2^+^ breast cancer tumors.

The ER+HER2-breast tumors have proven to be difficult to grow using the PDX model and other organoid cultures. We posit that the protocol ourlined here will provide a source of low passage primary ER+ breast tumor cells that could be useful in recreation of the original tumor environment in vitro.

This protocol provides the opportunity to obtain and sufficiently expand autologous BCC, CAF and TIL from breast tumor samples enabing their further study and characterization. Moreover, because each patient sample is unique, the derived cell types could be used to re-create the patient-specific tumor microenvironment *in vitro*. This, for example, could allow for the study of initial tumor response to therapy and the development of therapy resistance in a patient-specific manner. As well, the aulogous nature of the breast tumor cells obtained through this protocol enable studies aimed at gaining insight into mechanisms T cell dysfunction in breast tumors, thereby, help improve the current immunotherapy approaches for breast cancer patients.

