## Supplemental Figures and tables for "A robust protocol for the systematic collection and expansion of cells from ER^+^ breast cancer tumors and their matching tumor-adjacent tissues"

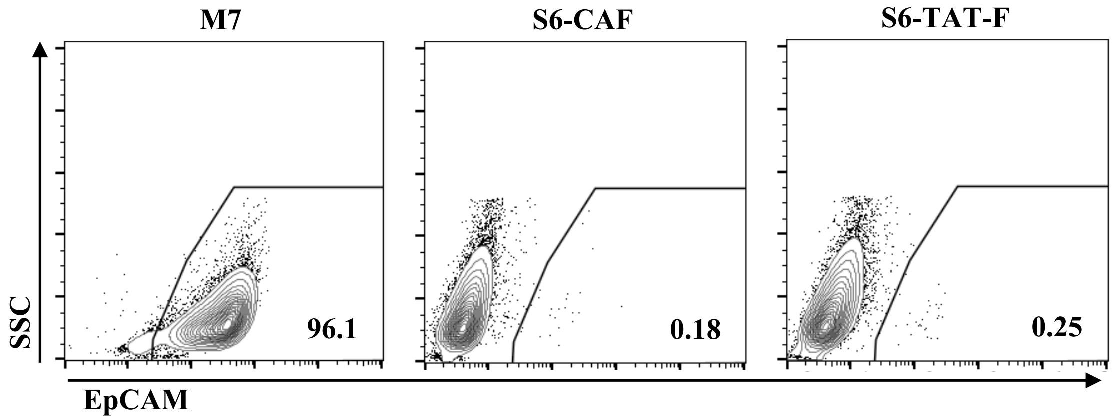


**Supplementary Figure 1. Supplementary Figure 1. In vitro expanded CAF and TAT-F cells contain few contaminating breast cancer cells.** In vitro expanded cancer-associated fibroblasts (CAF) and the tumor-adjacent tissue fibroblasts (TAT-F) were examined for the presence of contaminating epithelial cells adhesion molecule (EpCAM)-expressing breast cancer cells using fluorescent activated cell sorting (FACS). Representative FACS plots are shown for cells obtained from sample 6 (S6). The expression of EpCAM protein on the MCF7 (M7) was used as positive control. The expression of EpCAM on CAF and TAT-F developed in this study ranged from 0% to 0.4%. The data was analyzed using FloJo software Version 9.9.5.

| Receptor ID | M7 | MB-231 | S2 | S3 | S4 | S5 | N1 | M10A | N1-NAF | S4-CAF |
|---|---|---|---|---|---|---|---|---|---|---|
| CD1a | 2.34 | 0.287 | 0.177 | 0.315 | 0.053 | 0.164 | 0.71 | 0.251 | 0.258 | 0.741 |
| CD1b | 1.67 | 0.585 | 0.159 | 0.515 | 0.706 | 0.247 | 2.84 | 0.49 | 0.265 | 0.594 |
| CD1d | 1.94 | 0.633 | 0.129 | 0.986 | 0.237 | 0.227 | 1.28 | 0.35 | 0.297 | 0.285 |
| CD2 | 1.83 | 0.878 | 0.208 | 1 | 0.184 | 0.29 | 1.22 | 0.249 | 0.14 | 0.53 |
| CD3 | 1.93 | 0.515 | 0.149 | 0.818 | 0.758 | 0.187 | 1.4 | 0.08 | 0.318 | 0.377 |
| CD4 | 1.43 | 0.396 | 0.198 | 1.31 | 1.23 | 0.418 | 0.944 | 0.138 | 0.301 | 0.613 |
| CD4v4 | 1.49 | 1.03 | 0.258 | 1.2 | 0.362 | 0.332 | 0.789 | 0.528 | 0.402 | 0.497 |
| CD5 | 1.5 | 0.723 | 0.169 | 1.9 | 0.115 | 0.235 | 2.51 | 0.452 | 0.421 | 1.16 |
| CD6 | 1.61 | 0.357 | 0.129 | 2.03 | 0.581 | 0.196 | 0.909 | 0.208 | 0.432 | 0.713 |
| CD7 | 1.41 | 1.2 | 0.198 | 1.64 | 0.208 | 0.244 | 1.37 | 0.117 | 0.419 | 0.665 |
| CD8a | 1.2 | 0.303 | 0.07 | 1.25 | 0.265 | 0.232 | 0.296 | 0.201 | 0.512 | 0.654 |
| CD8b | 1.9 | 0.199 | 0.178 | 0.681 | 1.29 | 0.26 | 0.236 | 0.846 | 0.671 | 0.613 |
| CD9 | 30.3 | 49.7 | 70.2 | 85.8 | 55.7 | 64.1 | 9.96 | 81.6 | 44.8 | 12.6 |
| CD10 | 2.22 | 0.377 | 3.48 | 17.9 | 3.84 | 1.6 | 82.5 | 0.45 | 20.8 | 72.5 |
| CD11a | 1.98 | 0.457 | 0.596 | 0.802 | 0.62 | 0.421 | 0.451 | 0.548 | 0.603 | 0.494 |
| CD11b | 2.29 | 0.506 | 0.228 | 1.16 | 0.096 | 0.173 | 0.589 | 0.228 | 0.517 | 0.684 |
| CD11c | 1.85 | 0.288 | 0.109 | 0.808 | 0.447 | 0.617 | 0.651 | 0.102 | 1.69 | 2.01 |
| CD13 | 1.39 | 0.348 | 94 | 19.9 | 16.5 | 92.6 | 94.5 | 0.257 | 85.7 | 95.1 |
| CD14 | 1.81 | 0.516 | 1.05 | 3.47 | 0.447 | 0.192 | 0.388 | 0.234 | 0.776 | 0.504 |
| CD15 | 34.8 | 0.765 | 1.63 | 1.27 | 4.41 | 0.808 | 3.13 | 64.2 | 1.46 | 2.09 |
| CD15s | 1.44 | 0.387 | 0.179 | 2.33 | 10.2 | 2.63 | 0.228 | 0.226 | 0.313 | 0.784 |
| CD16 | 1.26 | 0.81 | 0.169 | 0.997 | 0.209 | 0.221 | 0.337 | 0.507 | 0.347 | 0.931 |
| CD18 | 1.08 | 0.584 | 0.129 | 0.539 | 0.18 | 0.218 | 0.401 | 0.236 | 0.415 | 0.897 |
| CD19 | 1.76 | 0.278 | 0.089 | 0.802 | 1.38 | 0.256 | 0.154 | 0.442 | 0.236 | 1.01 |
| CD20 | 1.89 | 0.318 | 0.099 | 0.575 | 0.545 | 0.128 | 0.207 | 0.189 | 0.45 | 0.565 |
| CD21 | 2.32 | 0.427 | 0.12 | 0.595 | 0.231 | 0.234 | 0.486 | 0.332 | 0.538 | 0.564 |
| CD22 | 1.73 | 0.278 | 0.139 | 0.738 | 0.175 | 0.118 | 0.194 | 0.38 | 0.456 | 0.552 |
| CD23 | 2.45 | 0.387 | 0.149 | 0.715 | 0.927 | 0.421 | 0.14 | 0.818 | 0.272 | 0.94 |
| CD24 | 9.28 | 0.645 | 83.4 | 76.3 | 81.9 | 40.5 | 0.426 | 94.7 | 0.376 | 0.559 |
| CD25 | 1.87 | 0.198 | 0.636 | 0.73 | 0.294 | 0.199 | 0.323 | 0.257 | 1.85 | 1.25 |
| CD26 | 1.42 | 0.525 | 2.12 | 0.685 | 0.608 | 1.16 | 56.7 | 0.74 | 82.3 | 90.7 |
| CD27 | 1.35 | 0.367 | 0.139 | 0.725 | 0.678 | 0.247 | 0.647 | 0.457 | 3.31 | 2.41 |
| CD28 | 1.54 | 0.775 | 0.119 | 0.95 | 0.346 | 0.167 | 0.412 | 0.268 | 2.04 | 2.35 |
| CD29 | 1.66 | 0.456 | 0.288 | 1.56 | 0.405 | 0.298 | 14 | 0.339 | 0.806 | 19 |
| CD30 | 1.46 | 0.515 | 0.129 | 0.635 | 0.249 | 0.141 | 0.194 | 0.338 | 0.746 | 1.11 |
| CD31 | 1.72 | 0.298 | 0.129 | 0.491 | 1.17 | 0.238 | 0.314 | 0.436 | 0.529 | 1.34 |
| CD32 | 1.67 | 0.278 | 0.109 | 0.484 | 0.538 | 0.159 | 0.271 | 0.371 | 0.31 | 0.885 |
| CD33 | 2.08 | 0.358 | 0.11 | 0.845 | 0.134 | 0.222 | 0.244 | 0.244 | 0.675 | 0.704 |
| CD34 | 1.73 | 0.367 | 0.139 | 0.54 | 0.201 | 0.12 | 0.277 | 0.208 | 0.291 | 0.563 |
| CD35 | 2.41 | 0.318 | 0.189 | 0.546 | 0.135 | 0.279 | 0.239 | 0.157 | 0.498 | 0.611 |
| CD36 | 1.94 | 0.387 | 0.189 | 0.574 | 0.235 | 0.176 | 1.34 | 0.941 | 13.5 | 2.83 |
| CD37 | 1.45 | 0.288 | 0.149 | 0.545 | 0.212 | 0.154 | 0.202 | 0.41 | 0.761 | 0.853 |
| CD38 | 1.52 | 0.268 | 0.1 | 0.469 | 0.286 | 0.093 | 0.386 | 0.34 | 0.736 | 0.817 |
| CD39 | 1.64 | 0.288 | 0.1 | 0.488 | 0.23 | 0.157 | 0.368 | 0.333 | 0.636 | 0.598 |
| CD40 | 1.84 | 0.367 | 0.348 | 0.83 | 0.296 | 0.317 | 1.69 | 0.158 | 0.225 | 0.846 |
| CD41a | 2.12 | 0.505 | 0.229 | 0.632 | 0.227 | 0.131 | 0.214 | 0.401 | 0.237 | 0.615 |
| CD41b | 1.39 | 0.386 | 0.229 | 0.939 | 0.224 | 0.11 | 0.279 | 0.515 | 0.354 | 0.758 |
| CD42a | 2.11 | 0.377 | 0.267 | 0.765 | 1.71 | 0.265 | 0.348 | 0.444 | 0.891 | 0.646 |
| CD42b | 2.04 | 0.347 | 0.119 | 0.594 | 0.403 | 0.137 | 0.238 | 0.479 | 0.444 | 0.872 |
| CD43 | 2.64 | 0.278 | 0.16 | 1.1 | 0.533 | 0.452 | 0.837 | 0.478 | 1.52 | 0.939 |
| CD44 | 1.72 | 99.1 | 97.2 | 90.8 | 95.1 | 97.2 | 96.7 | 71.9 | 92.7 | 95.9 |
| CD45 | 2.27 | 0.428 | 0.497 | 0.937 | 0.18 | 0.184 | 0.284 | 0.216 | 0.667 | 1.13 |
| CD45RA | 2.01 | 0.436 | 0.119 | 0.71 | 0.527 | 0.705 | 0.604 | 0.649 | 2.26 | 1.81 |
| CD45RB | 1.67 | 0.376 | 0.139 | 0.656 | 0.143 | 0.165 | 0.373 | 0.421 | 0.539 | 0.417 |
| CD45RO | 1.78 | 0.415 | 0.159 | 0.675 | 0.355 | 0.563 | 0.621 | 0.239 | 0.863 | 1.32 |
| CD46 | 15.7 | 95.7 | 90.8 | 81.5 | 60.2 | 77 | 91.1 | 93.3 | 54.8 | 88.5 |
| CD47 | 24.3 | 16.6 | 13.2 | 15.9 | 3.13 | 10.7 | 44.8 | 83.7 | 5.41 | 57.9 |
| CD48 | 2.25 | 0.513 | 0.209 | 1.09 | 0.489 | 0.776 | 0.504 | 0.736 | 6.71 | 1.79 |
| CD49a | 1.81 | 1.13 | 0.468 | 1.01 | 0.485 | 0.625 | 15 | 0.694 | 0.99 | 11.4 |
| CD49b | 20.7 | 78.1 | 41.4 | 65.6 | 67.5 | 26.3 | 73.6 | 50.5 | 47.8 | 89.5 |
| CD49c | 3.39 | 76.6 | 91.7 | 92.5 | 88.2 | 80.9 | 50.9 | 57.5 | 5.45 | 60.6 |
| CD49d | 3.01 | 0.592 | 0.377 | 1.75 | 1.27 | 0.332 | 22.1 | 0.349 | 5.2 | 38.5 |
| CD49e | 2.1 | 15.2 | 6.37 | 16.7 | 22.2 | 3.23 | 68.3 | 7.25 | 54.2 | 37.3 |
| CD50 | 1.98 | 0.513 | 0.179 | 0.669 | 0.29 | 0.212 | 0.505 | 0.2 | 0.747 | 1.47 |
| CD51/61 | 1.65 | 0.435 | 0.238 | 0.596 | 0.665 | 0.292 | 0.923 | 0.453 | 2.1 | 2.4 |
| CD53 | 1.9 | 0.403 | 0.109 | 0.837 | 0.261 | 0.293 | 0.377 | 0.27 | 0.636 | 0.874 |
| CD54 | 3.91 | 88.4 | 52.4 | 18.3 | 9.01 | 29.8 | 36.3 | 19.6 | 3.65 | 75 |
| CD55 | 21.3 | 97.1 | 87.9 | 68.5 | 47.8 | 79.8 | 94 | 83.2 | 47.1 | 71.1 |
| CD56 | 1.69 | 0.576 | 0.636 | 0.636 | 0.381 | 0.473 | 0.405 | 0.438 | 2.97 | 3.36 |
| CD57 | 2.36 | 0.484 | 0.568 | 1.28 | 0.871 | 1.55 | 0.908 | 1.11 | 2.84 | 2.09 |
| CD58 | 5.85 | 85.4 | 38 | 31.9 | 12 | 22.8 | 30.6 | 90.5 | 5.5 | 6.43 |
| CD59 | 39.8 | 86.6 | 76.5 | 99.5 | 98.4 | 95.4 | 92.4 | 96.5 | 95.1 | 97.6 |
| CD61 | 2.29 | 0.414 | 0.268 | 0.997 | 0.336 | 0.245 | 0.784 | 0.281 | 1.2 | 0.601 |
| CD62E | 3.34 | 0.582 | 0.477 | 1.12 | 0.773 | 0.756 | 0.803 | 0.792 | 2.6 | 5.34 |
| CD62L | 1.47 | 0.365 | 0.109 | 0.775 | 0.295 | 0.258 | 0.339 | 0.255 | 0.558 | 1.16 |
| CD62P | 1.71 | 0.385 | 0.16 | 0.805 | 0.248 | 0.274 | 0.906 | 0.319 | 1.45 | 1.05 |
| CD63 | 9.08 | 15.7 | 4.72 | 1.77 | 2.16 | 12.8 | 55.9 | 70.4 | 41.1 | 31.8 |
| CD64 | 1.62 | 0.335 | 0.249 | 0.553 | 0.235 | 0.62 | 0.457 | 0.562 | 0.687 | 1.23 |
| CD66 | 2.39 | 0.542 | 90.4 | 23.6 | 24.7 | 89.5 | 0.573 | 66 | 1.26 | 1.08 |
| CD66b | 1.31 | 0.295 | 0.616 | 1.08 | 0.267 | 0.253 | 0.175 | 0.439 | 1.04 | 1.34 |
| CD66f | 1.68 | 0.256 | 0.08 | 0.944 | 0.477 | 0.787 | 0.141 | 0.41 | 1.12 | 2.49 |
| CD69 | 2.01 | 0.463 | 0.119 | 0.696 | 0.206 | 0.187 | 0.467 | 0.258 | 0.851 | 69.2 |
| CD70 | 2.06 | 0.472 | 0.189 | 0.82 | 0.626 | 0.249 | 0.418 | 0.6 | 0.999 | 2.82 |
| CD71 | 23.2 | 95.5 | 3.17 | 36.9 | 44.7 | 49.2 | 79.7 | 97.2 | 47 | 71.5 |
| CD72 | 2.21 | 0.481 | 0.308 | 1.02 | 0.779 | 0.239 | 0.434 | 0.405 | 3.8 | 3.2 |
| CD73 | 2.54 | 56.5 | 4.5 | 92.3 | 65.5 | 3.04 | 88.1 | 1.51 | 77.9 | 85.1 |
| CD74 | 2.27 | 1.72 | 0.269 | 1.37 | 0.595 | 0.204 | 0.692 | 0.34 | 2.45 | 3.23 |
| CD75 | 5.64 | 1.17 | 0.587 | 1.56 | 0.784 | 0.345 | 0.864 | 1.8 | 3.88 | 4.24 |
| CD77 | 2.31 | 0.275 | 0.18 | 1.01 | 0.603 | 0.289 | 0.265 | 0.511 | 2.26 | 2.56 |
| CD79b | 1.92 | 0.708 | 0.169 | 0.735 | 0.339 | 0.284 | 0.688 | 0.427 | 1.54 | 1.84 |
| CD80 | 1.67 | 0.491 | 0.229 | 1.03 | 0.498 | 0.345 | 0.776 | 0.659 | 1.7 | 5.86 |
| CD81 | 21.3 | 37.2 | 45.6 | 24.9 | 16.7 | 31.9 | 91.1 | 87 | 85.3 | 91.4 |
| CD83 | 1.85 | 0.521 | 0.338 | 0.745 | 0.471 | 0.17 | 0.528 | 0.573 | 2.31 | 2.08 |
| CD84 | 1.97 | 0.588 | 0.149 | 0.877 | 0.369 | 0.477 | 0.816 | 0.5 | 2.69 | 2.73 |
| CD85 | 1.85 | 0.636 | 0.249 | 1.2 | 0.245 | 0.175 | 0.39 | 0.17 | 2.79 | 2.22 |
| CD86 | 1.58 | 0.365 | 0.069 | 0.296 | 0.106 | 0.114 | 0.378 | 0.171 | 0.442 | 0.613 |
| CD87 | 1.77 | 0.285 | 0.09 | 0.585 | 0.154 | 0.107 | 0.557 | 0.191 | 0.935 | 0.721 |
| CD88 | 1.5 | 0.404 | 0.079 | 0.704 | 0.093 | 0.105 | 0.742 | 0.127 | 0.525 | 0.476 |
| CD89 | 1.31 | 0.609 | 0.079 | 0.572 | 0.232 | 0.115 | 0.639 | 0.115 | 1.67 | 1.51 |
| CD90 | 1.73 | 0.276 | 1.05 | 0.748 | 1.65 | 1.05 | 95 | 0.264 | 80.7 | 91.6 |
| CD91 | 2.15 | 1.33 | 0.267 | 0.708 | 0.557 | 0.316 | 66.1 | 1.75 | 9.06 | 31.8 |
| CDw93 | 1.46 | 0.462 | 0.089 | 0.768 | 0.135 | 0.259 | 0.587 | 0.329 | 2.56 | 1.33 |
| CD94 | 1.9 | 0.856 | 0.109 | 0.431 | 0.076 | 0.158 | 0.619 | 0.125 | 1.2 | 1.38 |
| CD95 | 2.03 | 0.895 | 2.17 | 5.03 | 1.05 | 0.547 | 19.5 | 1.73 | 7.91 | 65.2 |
| CD97 | 2.36 | 68.5 | 11.1 | 0.946 | 2.41 | 9.26 | 40.5 | 1.15 | 11.5 | 2.72 |
| CD98 | 50.6 | 30.2 | 68.2 | 97.8 | 87.4 | 90.2 | 37.4 | 92.5 | 9.95 | 44.1 |
| CD99 | 19.3 | 34.3 | 4.64 | 27.8 | 4.12 | 0.816 | 76.5 | 23 | 66.5 | 92.9 |
| CD99R | 1.52 | 0.59 | 0.119 | 0.701 | 0.15 | 0.139 | 16.4 | 0.143 | 9.01 | 58.4 |
| CD100 | 1.62 | 0.305 | 0.13 | 0.656 | 0.125 | 0.171 | 0.268 | 0.247 | 1.71 | 1.71 |
| CD102 | 1.75 | 1.1 | 0.05 | 1.15 | 0.058 | 0.225 | 0.216 | 0.093 | 0.493 | 1.2 |
| CD103 | 1.55 | 0.296 | 0.079 | 1.22 | 0.181 | 0.173 | 0.234 | 0.175 | 0.466 | 0.34 |
| CD105 | 1.52 | 0.237 | 0.129 | 0.575 | 0.196 | 0.243 | 7.27 | 0.221 | 0.669 | 12.4 |
| CD106 | 2.07 | 0.335 | 0.199 | 0.512 | 0.327 | 0.156 | 14.5 | 0.253 | 0.442 | 0.706 |
| CD107a | 7.05 | 4.04 | 0.377 | 82.8 | 0.257 | 0.309 | 0.806 | 0.502 | 4.18 | 9.37 |
| CD107b | 7.01 | 5.88 | 0.279 | 32 | 0.275 | 0.229 | 0.295 | 0.708 | 1.54 | 8.06 |
| CD108 | 3.04 | 12.9 | 0.09 | 4.27 | 0.219 | 0.171 | 0.565 | 2.32 | 9.87 | 32.3 |
| CD109 | 2.59 | 0.384 | 0.099 | 1.68 | 0.149 | 0.192 | 20.9 | 1.59 | 1.16 | 3.33 |
| CD112 | 1.73 | 0.216 | 0.129 | 0.543 | 0.169 | 0.146 | 0.328 | 0.32 | 0.76 | 0.798 |
| CD114 | 1.41 | 0.226 | 0.06 | 0.564 | 0.597 | 0.196 | 0.346 | 0.23 | 0.74 | 0.95 |
| CD116 | 1.56 | 0.286 | 0.06 | 0.534 | 0.255 | 0.203 | 0.18 | 0.249 | 0.348 | 0.683 |
| CD117 | 1.79 | 0.296 | 7.49 | 0.675 | 0.112 | 1.93 | 0.157 | 0.122 | 0.313 | 0.871 |
| CD118 | 1.75 | 0.395 | 0.08 | 0.476 | 0.109 | 0.195 | 0.215 | 0.232 | 0.366 | 0.775 |
| CD119 | 2.11 | 0.315 | 0.08 | 0.853 | 0.227 | 0.259 | 0.156 | 0.215 | 0.276 | 0.433 |
| CD120a | 2.3 | 0.533 | 0.07 | 0.786 | 0.165 | 0.109 | 0.209 | 0.212 | 0.307 | 0.787 |
| CD121a | 2.13 | 0.384 | 0.05 | 0.383 | 0.297 | 0.14 | 0.33 | 0.204 | 0.424 | 0.825 |
| CD121b | 2.57 | 0.326 | 0.05 | 0.445 | 0.08 | 0.147 | 0.032 | 0.156 | 0.708 | 0.635 |
| CD122 | 2.28 | 0.71 | 0.07 | 0.542 | 0.169 | 0.154 | 0.161 | 0.379 | 0.472 | 0.438 |
| CD123 | 3 | 0.474 | 0.119 | 0.461 | 0.125 | 0.161 | 0.225 | 0.228 | 0.457 | 0.475 |
| CD124 | 2.67 | 0.306 | 0.11 | 0.489 | 0.099 | 0.19 | 0.292 | 0.097 | 0.837 | 0.76 |
| CD126 | 1.75 | 38.4 | 0.13 | 0.519 | 0.497 | 0.196 | 7.9 | 4.86 | 25.6 | 61.3 |
| CD127 | 1.85 | 0.276 | 0.07 | 0.478 | 0.685 | 0.209 | 0.214 | 0.337 | 0.991 | 0.978 |
| CD128b | 1.68 | 1.32 | 0.05 | 3.83 | 0.118 | 0.142 | 0.295 | 0.213 | 0.213 | 0.674 |
| CD130 | 1.64 | 0.482 | 0.04 | 0.993 | 0.417 | 0.141 | 0.403 | 0.37 | 0.46 | 1.03 |
| CD134 | 1.52 | 0.237 | 0.07 | 0.411 | 0.228 | 0.12 | 0.444 | 0.304 | 0.305 | 0.822 |
| CD135 | 2.03 | 0.314 | 0.104 | 0.296 | 0.096 | 0.137 | 0.1 | 0.211 | 0.33 | 0.402 |
| CD137 | 2.25 | 0.246 | 0.119 | 0.419 | 0.117 | 0.153 | 0.179 | 0.062 | 0.293 | 0.772 |
| CD137L | 2.43 | 0.325 | 0.04 | 0.524 | 0.251 | 0.164 | 0.115 | 0.448 | 0.189 | 0.632 |
| CD138 | 2.93 | 0.364 | 0.139 | 1.1 | 9.85 | 1.16 | 0.425 | 30.9 | 0.916 | 12.9 |
| CD140a | 2.29 | 0.637 | 0.15 | 0.397 | 0.163 | 0.178 | 16.4 | 0.348 | 2.98 | 14 |
| CD140b | 2.77 | 0.325 | 0.08 | 0.935 | 0.571 | 0.229 | 77.1 | 0.287 | 8.71 | 15.4 |
| CD141 | 2.91 | 10 | 0.149 | 35.7 | 23.8 | 0.506 | 0.706 | 8.07 | 1.43 | 0.98 |
| CD142 | 1.33 | 81.5 | 2.42 | 46.7 | 67.7 | 3 | 5.65 | 0.368 | 7.81 | 40.3 |
| CD144 | 1.66 | 0.296 | 0.06 | 0.692 | 1.07 | 0.107 | 0.581 | 0.357 | 0.551 | 0.938 |
| CD146 | 3.5 | 43.8 | 74.1 | 11.2 | 12.8 | 60.2 | 1.01 | 0.538 | 1.6 | 48.9 |
| CD147 | 28 | 88 | 92.7 | 97.3 | 92.7 | 89.5 | 94.6 | 83.8 | 67.6 | 91.1 |
| CD150 | 1.73 | 0.314 | 0.639 | 1 | 0.184 | 0.434 | 0.401 | 0.334 | 1.3 | 1.55 |
| CD151 | 11.1 | 16.4 | 31.9 | 1.19 | 6.12 | 18 | 80.3 | 60.5 | 77 | 86.7 |
| CD152 | 2.16 | 0.227 | 0.23 | 1.31 | 0.062 | 0.073 | 0.429 | 0.28 | 0.722 | 0.923 |
| CD153 | 2.15 | 0.285 | 0.279 | 1.42 | 0.234 | 0.299 | 0.551 | 0.511 | 1.72 | 1.35 |
| CD154 | 2.4 | 0.314 | 0.09 | 0.778 | 0.161 | 0.153 | 0.419 | 0.309 | 0.322 | 1.43 |
| CD158a | 2.38 | 0.344 | 0.1 | 0.988 | 0.22 | 0.126 | 0.258 | 0.302 | 0.971 | 0.928 |
| CD158b | 2.68 | 0.246 | 0.199 | 1.02 | 0.179 | 0.124 | 0.186 | 0.277 | 0.529 | 0.948 |
| CD161 | 1.82 | 1.48 | 0.09 | 1.06 | 0.245 | 0.088 | 0.381 | 0.276 | 0.472 | 1.29 |
| CD162 | 1.52 | 0.353 | 0.11 | 1.07 | 0.241 | 0.258 | 0.427 | 0.142 | 0.473 | 1.21 |
| CD163 | 1.62 | 0.304 | 0.09 | 1.15 | 0.377 | 0.139 | 0.31 | 0.294 | 0.858 | 1.61 |
| CD164 | 36.4 | 92.1 | 4.17 | 24.6 | 8.65 | 3.37 | 13.5 | 54.6 | 75 | 39.4 |
| CD165 | 1.91 | 0.235 | 0.09 | 0.982 | 0.235 | 0.121 | 0.934 | 0.405 | 1.68 | 2.64 |
| CD166 | 13.9 | 40.2 | 6.42 | 12.3 | 6.58 | 2.35 | 18.5 | 49.8 | 59.2 | 72.6 |
| CD171 | 8.15 | 7.75 | 47.8 | 1.08 | 1.26 | 2.97 | 0.195 | 49.2 | 0.664 | 1.14 |
| CD172b | 1.85 | 0.276 | 0.378 | 1.26 | 0.221 | 0.133 | 0.572 | 0.472 | 0.777 | 1.31 |
| CD177 | 1.86 | 0.235 | 0.319 | 0.713 | 0.146 | 0.17 | 0.446 | 0.48 | 0.269 | 0.782 |
| CD178 | 2.08 | 0.295 | 0.169 | 1.04 | 0.208 | 0.169 | 0.444 | 0.141 | 0.38 | 0.768 |
| CD180 | 2.18 | 0.343 | 0.1 | 1.52 | 0.248 | 0.056 | 0.261 | 0.384 | 0.917 | 1.02 |
| CD181 | 2.36 | 0.44 | 0.179 | 1.19 | 0.189 | 0.151 | 0.312 | 0.214 | 0.27 | 0.74 |
| CD183 | 1.85 | 0.295 | 0.11 | 1.14 | 0.199 | 0.218 | 0.338 | 0.155 | 0.601 | 1.67 |
| CD184 | 1.43 | 2.23 | 0.16 | 1.59 | 0.212 | 0.217 | 0.286 | 3.4 | 0.294 | 1.88 |
| CD193 | 1.63 | 0.361 | 0.119 | 1.71 | 0.52 | 0.1 | 0.358 | 0.485 | 1.93 | 1.74 |
| CD195 | 1.42 | 0.353 | 0.04 | 1.59 | 0.241 | 0.303 | 0.139 | 0.111 | 1.41 | 1.16 |
| CD196 | 1.93 | 0.263 | 0.14 | 1.24 | 0.337 | 0.084 | 0.289 | 0.349 | 0.741 | 1.71 |
| CD197 | 2.47 | 0.285 | 0.11 | 1.74 | 0.197 | 0.101 | 0.561 | 0.23 | 0.485 | 1.7 |
| CD200 | 1.66 | 0.392 | 0.377 | 1.11 | 0.215 | 0.331 | 0.271 | 0.316 | 1.1 | 1.87 |
| CD205 | 1.8 | 0.206 | 53.2 | 2.91 | 5.15 | 32.6 | 0.901 | 0.554 | 1.04 | 1.76 |
| CD206 | 1.89 | 0.333 | 0.089 | 1.24 | 0.316 | 0.197 | 0.238 | 0.287 | 0.821 | 1.03 |
| CD209 | 2.61 | 0.939 | 0.628 | 2.23 | 0.705 | 0.632 | 2.09 | 0.564 | 1.58 | 4.99 |
| CD220 | 1.95 | 0.264 | 0.13 | 1.71 | 0.261 | 0.228 | 0.459 | 0.569 | 1.2 | 1.12 |
| CD221 | 2.68 | 0.186 | 0.259 | 1.37 | 0.161 | 0.158 | 0.47 | 0.717 | 0.719 | 1.96 |
| CD226 | 2.16 | 0.323 | 0.13 | 1.24 | 0.241 | 0.596 | 0.301 | 0.528 | 0.688 | 2.62 |
| CD227 | 18.7 | 7.33 | 23.4 | 3.03 | 11.9 | 40.7 | 4.08 | 29.7 | 4.95 | 3.24 |
| CD229 | 2.16 | 0.554 | 0.149 | 1.9 | 0.578 | 0.405 | 0.463 | 0.531 | 3.52 | 4.97 |
| CD231 | 1.52 | 0.35 | 0.089 | 1.86 | 0.305 | 0.276 | 0.307 | 0.521 | 2.41 | 3.02 |
| CD235a | 2.11 | 0.458 | 0.11 | 0.94 | 0.236 | 0.118 | 0.409 | 0.68 | 1.81 | 2.77 |
| CD243 | 1.83 | 0.488 | 0.22 | 1.78 | 0.327 | 0.141 | 0.529 | 0.307 | 1.86 | 3.82 |
| CD244 | 1.67 | 0.321 | 0.189 | 1.51 | 0.34 | 0.421 | 0.135 | 0.12 | 1.98 | 2.45 |
| CD255 | 1.68 | 0.253 | 0.12 | 1.26 | 0.16 | 0.171 | 0.405 | 0.475 | 1.98 | 4.63 |
| CD268 | 1.47 | 0.332 | 0.14 | 1.34 | 0.363 | 0.161 | 0.404 | 0.28 | 1.16 | 2.42 |
| CD271 | 2 | 9.35 | 1.42 | 38.8 | 6.83 | 2.72 | 0.556 | 0.31 | 1.71 | 1.46 |
| CD273 | 1.69 | 0.524 | 0.319 | 1.59 | 0.165 | 0.139 | 0.349 | 0.383 | 2.52 | 3.45 |
| CD274 | 2 | 0.585 | 0.249 | 1.88 | 0.233 | 0.184 | 0.539 | 0.322 | 1.95 | 3.04 |
| CD275 | 1.66 | 0.407 | 0.08 | 1.86 | 0.333 | 0.166 | 0.468 | 0.403 | 2.44 | 2.63 |
| CD278 | 2 | 0.281 | 0.12 | 1.68 | 0.633 | 0.199 | 0.723 | 0.174 | 1.87 | 3.08 |
| CD279 | 1.46 | 0.308 | 1.07 | 0.404 | 0.151 | 0.399 | 1.91 | 0.206 | 0.281 | 0.733 |
| CD282 | 1.26 | 0.378 | 0.893 | 1.61 | 0.448 | 0.408 | 0.691 | 0.5 | 0.285 | 0.35 |
| CD305 | 1.57 | 0.179 | 0.222 | 0.619 | 0.173 | 0.409 | 2.17 | 0.364 | 0.326 | 1 |
| D309 | 1.49 | 0.377 | 0.13 | 1.06 | 1.2 | 0.351 | 0.161 | 0.235 | 0.391 | 0.474 |
| CD314 | 1.76 | 0.238 | 0.1 | 0.588 | 0.714 | 0.253 | 0.755 | 0.741 | 0.302 | 0.982 |
| CD321 | 41.8 | 0.685 | 22.7 | 32.2 | 77.9 | 18.3 | 0.837 | 71.4 | 0.155 | 0.755 |
| CDw327 | 2 | 0.397 | 1.36 | 0.276 | 0.261 | 0.459 | 0.345 | 0.373 | 0.717 | 0.773 |
| CDw328 | 1.69 | 0.457 | 0.351 | 0.519 | 0.431 | 0.459 | 0.415 | 0.728 | 0.82 | 0.797 |
| CDw329 | 1.49 | 0.328 | 0.401 | 10.3 | 0.608 | 0.312 | 1.11 | 0.413 | 0.346 | 1.12 |
| CD335 | 1.71 | 0.308 | 0.08 | 15.5 | 0.221 | 0.509 | 0.981 | 1 | 0.582 | 0.46 |
| CD336 | 0.6 | 0.308 | 0.08 | 0.378 | 0.121 | 0.684 | 1.11 | 0.554 | 0.498 | 0.535 |
| CD337 | 1.46 | 0.407 | 0.451 | 0.411 | 0.269 | 0.312 | 0.289 | 0.545 | 0.326 | 0.636 |
| CD338 | 5.36 | 0.317 | 1.17 | 0.641 | 0.134 | 0.245 | 0.749 | 13.7 | 0.417 | 1.02 |
| CD340 | 10.6 | 0.417 | 9.68 | 2.9 | 2.16 | 3.48 | 0.614 | 26 | 0.22 | 0.852 |
| abTCR | 2.28 | 0.357 | 0.382 | 1.7 | 0.952 | 0.804 | 0.682 | 0.156 | 0.144 | 1.28 |
| B2-uGlob | 16.9 | 93.2 | 90.2 | 83 | 39.6 | 80.7 | 82.1 | 50.6 | 80.2 | 85 |
| BLTR-1 | 1.31 | 0.328 | 0.671 | 0.325 | 0.171 | 0.176 | 0.344 | 0.244 | 0.147 | 0.724 |
| CLIP | 1.64 | 0.278 | 0.07 | 0.676 | 0.305 | 0.819 | 0.661 | 0.435 | 0.371 | 1.8 |
| CMRF-44 | 1.68 | 0.407 | 0.181 | 0.621 | 0.191 | 0.127 | 0.196 | 0.287 | 0.669 | 0.717 |
| CMRF-56 | 2.4 | 0.328 | 0.22 | 0.47 | 0.128 | 0.284 | 0.347 | 0.225 | 0.364 | 1.01 |
| EGF-r | 1.66 | 86.4 | 18 | 91.8 | 22.3 | 1.26 | 1.25 | 0.61 | 24.9 | 9.22 |
| Fmlp-r | 1.44 | 0.358 | 0.21 | 0.605 | 0.105 | 0.234 | 0.67 | 0.415 | 0.364 | 1.02 |
| gd TCR | 1.16 | 0.299 | 0.06 | 0.562 | 0.219 | 0.343 | 0.577 | 0.319 | 0.398 | 0.613 |
| CD372 | 1.74 | 0.308 | 9.17 | 0.484 | 0.774 | 2.95 | 1.62 | 0.358 | 0.907 | 0.331 |
| HLA-A,B,C | 36.6 | 97.9 | 85.1 | 92.1 | 56.5 | 79.4 | 92.5 | 72.3 | 87.5 | 93.3 |
| HLA-A2 | 38.9 | 99 | 96.8 | 0.539 | 0.042 | 0.255 | 92.5 | 92.1 | 0.49 | 92.6 |
| HLA-DQ | 3.49 | 0.843 | 0.565 | 1.09 | 0.358 | 0.559 | 0.904 | 0.46 | 0.386 | 1.2 |
| HLA-DR | 2.02 | 42.7 | 1.45 | 0.572 | 0.724 | 0.451 | 0.546 | 0.95 | 0.383 | 3.07 |
| HLA-DR,P,O | 1.65 | 47.7 | 2.81 | 0.626 | 0.792 | 0.547 | 0.22 | 0.454 | 0.271 | 0.618 |
| GD2 | 1.96 | 0.297 | 0.2 | 0.639 | 0.199 | 1.4 | 14.7 | 3.89 | 11.6 | 0.746 |
| MIC A/B | 1.82 | 4.06 | 6.29 | 0.362 | 4.62 | 2.98 | 7.43 | 14.7 | 0.802 | 6.84 |
| NKB1 | 2.01 | 0.437 | 0.11 | 0.448 | 0.256 | 0.128 | 0.328 | 0.507 | 0.335 | 0.97 |
| SSEA-1 | 35.4 | 0.668 | 1.85 | 0.979 | 5.22 | 0.256 | 0.863 | 73.5 | 0.501 | 1.39 |
| SSEA-4 | 6.02 | 84.9 | 0.13 | 0.329 | 0.252 | 0.157 | 0.892 | 83.4 | 14.6 | 73.9 |
| TRA-1-60 | 1.67 | 0.496 | 0.1 | 0.565 | 0.179 | 0.137 | 0.364 | 0.838 | 0.501 | 1.26 |
| TRA-1-81 | 1.31 | 0.386 | 0.209 | 0.516 | 0.17 | 0.167 | 0.387 | 0.383 | 0.221 | 0.714 |
| Vb 23 | 1.53 | 0.189 | 0.101 | 0.232 | 0.179 | 0.177 | 0.182 | 0.336 | 0.411 | 0.833 |
| Vb 8 | 1.63 | 0.278 | 0.12 | 0.367 | 0.234 | 0.422 | 0.441 | 0.341 | 0.336 | 0.476 |
| CD49f | 2.84 | 97.7 | 88.4 | 98.7 | 90 | 98.4 | 1.33 | 19.4 | 1.330 | 4.98 |
| CD104 | 2.95 | 37.3 | 85.9 | 96.9 | 89.7 | 90.5 | 0.921 | 6.06 | 0.921 | 2.17 |
| CD120b | 1.51 | 1.06 | 1.14 | 0.744 | 2.33 | 1.09 | 0.368 | 1.01 | 0.368 | 1.4 |
| CD132 | 1.64 | 0.744 | 0.743 | 1.02 | 2.09 | 1.21 | 0.697 | 0.78 | 0.697 | 0.824 |
| CD201 | 2.5 | 1.03 | 0.571 | 2.48 | 9.41 | 0.688 | 41.2 | 1.1 | 41.200 | 9.07 |
| CD210 | 2.03 | 0.504 | 0.1 | 0.773 | 1.86 | 0.508 | 0.33 | 1.21 | 0.330 | 1.21 |
| CD212 | 1.88 | 0.574 | 0.11 | 0.612 | 1.43 | 0.362 | 0.52 | 0.836 | 0.520 | 1.28 |
| CD267 | 1.85 | 0.535 | 0.18 | 0.719 | 1.43 | 0.339 | 0.194 | 0.938 | 0.194 | 1.91 |
| CD294 | 1.92 | 0.535 | 0.13 | 0.844 | 1.6 | 0.637 | 0.377 | 0.639 | 0.377 | 1.13 |
| CD326 | 43.8 | 0.993 | 76.7 | 5.75 | 19.2 | 53.3 | 0.292 | 93 | 0.292 | 0.374 |
| CLA | 2.14 | 0.654 | 0.19 | 32.1 | 12.6 | 3.68 | 0.526 | 0.897 | 0.526 | 1.13 |
| INT B7 | 2.07 | 0.602 | 0.2 | 1.37 | 1.38 | 0.51 | 0.457 | 1.58 | 0.457 | 1.5 |
| SSEA-3 | 3.52 | 0.417 | 0.16 | 0.444 | 1.7 | 0.46 | 0.311 | 2.21 | 0.311 | 1.47 |

**Supplementary Table 1. Frequency of *in vitro* expanded cells expressing different surface markers.** *In vitro* expanded breast cancer cells (BCC), cancer-associated fibroblasts (CAF), normal breast epithelial (N), and normal-associated fibroblasts (NAF) were used in a cell surface receptor antibody array using a flow cytometer. MCF7 (M7), MCF10A (M10A), and MDA-MB-231 (MB-231) cells were also examined for their expression of the 243 cell surface markers. The data was analyzed using FlowJo Version 9.9.5. The frequency of cells positive for the expression of each marker is included in the table and was used to generate a hierarchical cluster of the samples (**Fig. 2E**).
